# Tracking *kdr* Alleles Associated with Pyrethroid Resistance in *Aedes albopictus* across Italy: A Nationwide Genotypic Dataset by MosqIRIT Network

**DOI:** 10.64898/2026.06.08.730819

**Authors:** Carlo Maria De Marco, Verena Pichler, Federica Gobbo, Sara Manzi, Eleonora Rosso, Federica Toniolo, Elena Carra, Angelica Petrella, Annalisa Grisendi, Francesco Defilippo, Carlotta Tessarolo, Elisabetta Ercole, Annalisa Accorsi, Andrea Mosca, Filippo Cassina, Marco Di Domenico, Valeria Di Lollo, Matteo De Ascentis, Silvio Gerardo D’Alessio, Ilaria Congiu, Valentina Donati, Valeria Carioti, Fahimeh Bandieinia, Stefano Gavaudan, Cristina Canonico, Guido Favia, Francesca Racciatti, Roberta Spaccapelo, Celine Alami, Claudio de Martinis, Alessia Pucciarelli, Gerardo Picazio, Maurizio Viscardi, Loredana Capozzi, Maria Grazia Cariglia, Luana Violante, Cipriano Foxi, Daniele Dedola, Valentina Sini, Luca Ruiu, Alessia Vinci, Silvia Scibetta, Eugenia Oliveri, Stefano Reale, Maria Liliana Di Pasquale, Sara Villari, Beniamino Caputo, Alessandra della Torre

## Abstract

This data paper presents a curated, georeferenced dataset of the frequencies of the two main target site mutations (V1016G and F1534C) associated with resistance to pyrethroid insecticides in *Aedes albopictus* in Italy. Populations were collected in 102 out 107 Italian provinces between 2023 and 2025. Specimens were sampled by members of the Mosquito Insecticide Resistance Italian Network (MosqIRIT) as part of RN2 activities within the INF-ACT project. Genotyping was performed on 3,517 individuals by specific allele-specific PCR assays. Each record includes metadata on sampling site, administrative location, developmental stage, collection method, and mutation-specific genotype frequencies. To support spatial analysis modelling effort, the dataset integrates geographic, eco-climatic, and demographic data. This resource will support mosquito control programs, pyrethroid resistance monitoring and managing, as well as ecological modelling, and is compliant with the FAIR data program.

## CONTEXT

*Aedes albopictus*, commonly known as the Asian tiger mosquito, has become a widespread species of public health concern in Italy and across Europe due to its aggressive biting behaviour, adaptability to urban environments, and competence in transmitting exotic arboviruses such as dengue, chikungunya, and Zika (Schaffner et al., 2013; De Marco & Schaffner, 2026). In the last years, increasing numbers of dengue cases have been reported in Europe, where Italy experienced the largest outbreaks with 82 and 213 human cases in 2023 and 2024, respectively (ECDC, 2025a). In 2025, hundreds of chikungunya human cases have been reported both in France (>480) and Italy (>200) (ECDC, 2025b). To reduce mosquito nuisance and the risk of disease outbreaks, European guidelines for the surveillance of invasive mosquitoes recommend larval source reduction and larvicide applications (ECDC, 2012). In contrast, pyrethroid-based adulticidal interventions are recommended only in the cases of ongoing - or high risk - of virus transmission, when a fast and effective abatement of adult mosquitoes is necessary. Pyrethroids interact with the voltage-sensitive sodium channel (VSSC) and interfere with the transmission of nervous signals, resulting in fast knockdown and eventually death of the mosquito. However, first evidence of resistance to pyrethroids have been reported across Europe (Pichler et al., 2018; ECDC, 2023).

Pyrethroids are the main insecticide class for mosquito adulticide spraying registered in Europe for mosquito control, therefore, the development of resistance to pyrethroids could have a large impact. Pyrethroid resistance can arise due to either metabolic mechanisms (consisting in the over-expression of detoxification enzymes) or target site resistance (Hemingway et al., 2004) involving non-synonymous mutations (often referred to as knock-down resistance -kdr- mutations) in the target site of pyrethroids i.e. the VGSC. Kdr mutations impair the insecticide’s ability to bind and act on the VGSC, lowering thus the pyrethroids’ ability to induce paralysis and death. Monitoring the frequency of kdr mutations strongly associated with resistant phenotypes (Pichler et al., 2019) can be helpful for planning mosquito control strategies: detecting kdr mutations circulating at low frequency can help to set-up resistance management strategies before the mutation reaches concerningly high frequencies; in addition the knowledge on the kdr frequency in a given population can be useful for forecasting pyrethroid efficacy and choosing the best and most effective alternative control strategies (Uemura et al., 2024).

In *Ae. albopictus* the substitutions with strongest support for their role in conferring pyrethroid resistance are mutations V1016G and F1534C (Kasai et al., 2019; Moyes et al., 2017). V1016G has been reported across Europe, with highest frequencies in Italy (Pichler et al., 2019; 2021; 2022). On the other hand, the distribution of the F1534C mutation is geographically constrained to Eastern Europe, yet it exhibits exceptionally high allelic frequencies in Cyprus (84%) and Greece (45%) (Pichler et al. 2025).

We developed a harmonized, geo-referenced dataset of *Ae. albopictus* V1016G and F1534C frequencies in 102 out of 107 Italian provinces between 2023 and 2025. Specimens were sampled by members of the Mosquito Insecticide Resistance Italian Network (MosqIRIN) as part of RN2 activities within the INF-ACT project on “One health basic and translational research actions addressing unmet needs on emerging infectious diseases” (https://www.inf-act.it/). By combining high-resolution spatial data with standardized genotyping and environmental context, the dataset not only is compliant with the FAIR data program, but is expected to support downstream applications aimed at improving mosquito control strategies, pyrethroid resistance monitoring and managing, and advancing research on the ecology and evolution of insecticide resistance in *Ae. albopictus*.

## METHODS

### Mosquito Collection and V1016G and F1534C genotyping

*Aedes albopictus* specimens were sampled between 2023 and 2025 by either ovitraps, larval sampling or adult trapping with the aim to provide a baseline dataset of V1016G and F1534C frequencies in samples ≥30 individuals for each province in Italy. Larvae collected in the field or hatched from ovitrap-collected eggs were reared to adults under standard insectary conditions. No more than two adults emerged from a each ovitrap were analysed to avoid risk of consanguinity.

Genomic DNA was extracted from single specimens, using different manual extraction methods, DNA extraction kits (NZY Tissue gDNA Isolation kit – Nzytech, Portugal; DNeasy Blood and Tissue, Qiagen, Germany; Maxwell RSC Tissue DNA Kit, Promega, USA) or kit (BioSprint® 96 One-For-All Vet, Qiagen, Germany) in combination with semi-automatic extraction system (King-Fisher, Thermo Fisher Scientific, USA), according to the manufacturer’s instructions. Extracted DNA was stored at -20ºC until further analysis.

Genotyping was performed by each collaborating Institution by standardized AS-PCR methods (Pichler et al., 2021 for V1016G mutation, and Zhu et al., 2019 for F1534C mutation). Sequencing of a fragment surrounding F1534C mutation was performed, when possible, for specimens PCR-genotyped as heterozygous or homozygous for the 1534C allele using outer primers described by Zhu et al. (2019) and send for Sanger-sequencing at Eurofins Genomics (Ebersberg, Germany). Sequences were read using the software Chromas version 2.6.6 (Technelysium 2025).

### Data geo-referencing

To enable the use of the dataset in geospatial analyses while safeguarding the privacy of sampling locations (whose GPS coordinates are not included in the dataset), all mosquito sampling sites were spatially masked using a regular grid of 0.025 degrees in both latitude and longitude (corresponding to approximately 4 km^2^). In cases where coordinates were unavailable, site locations were manually geocoded using at least two online gazetteers (e.g., Google Maps and OpenStreetMap), ensuring cross-validation of place names and administrative boundaries. When multiple samples fell within the same grid cell, genotyping results for the *kdr* assay were aggregated accordingly. For province-level summaries, allele frequencies, spatial joins were performed using the GADM administrative boundary dataset (GADM, 2024).

### Environmental and demographic data integration

To support future ecological modelling and spatial analyses, each mosquito sampling site was enriched with standardized environmental and demographic covariates derived from publicly available geospatial datasets. Specifically:

<u>Land cover data</u> were obtained from the CORINE Land Cover 2018 (CLC18) dataset, produced by the Copernicus Land Monitoring Service. Using a 300-meter buffer around each sampling coordinate, the proportion of land cover classes (e.g., urban fabric, agricultural areas, forests) was calculated. Each site record includes both the categorical CLC18 class with the highest coverage and the percentage distribution of all classes within the buffer (ISPRA, 2018).

<u>Air temperature data</u> were extracted from the ERA5 reanalysis product provided by the Copernicus Climate Data Store (Muñoz Sabater, 2019). For each site, the mean 2-meter air temperature (°C) was computed for the full calendar year 2024, using a monthly temporal resolution and spatial interpolation at 0.25 in both latitude and longitude. Temperature values were matched to each sampling location based on the nearest grid cell centroid.

<u>Population data</u> were derived from the Italian National Institute of Statistics (ISTAT) 2021 census estimates (ISTAT, 2021). Using a 300-meter circular buffer around each sampling point, population counts were extracted from regional demographic layers to quantify human density in the immediate vicinity of each mosquito collection site. This provides a proxy for anthropogenic pressure and potential insecticide exposure.

All spatial extractions and buffer operations were performed in QGIS v3.32 and Python using the geopandas library v1.1.1. (Bossche et al., 2025). Spatial covariates were harmonized to the WGS 84 coordinate reference system (EPSG:4326) to ensure compatibility with the rest of the dataset. Source code is available in GitHub (De Marco, 2026).

## RESULTS

Overall, 3,517 and 2,717 *Ae. albopictus* specimens from 102 of 107 Italian provinces were successfully genotyped for V1016G and F1534C mutations, respectively. Table 1 summarises the number and *kdr*-genotypes of individuals analysed and the *kdr*-allele frequencies in each province. The median sample size/province was 30. Figure 1 shows V1016G and F1534C mutation frequencies in the 198 sampling sites with ≥ 5 individuals (median size number per site = 14 and 11, respectively).

**Table 1.**
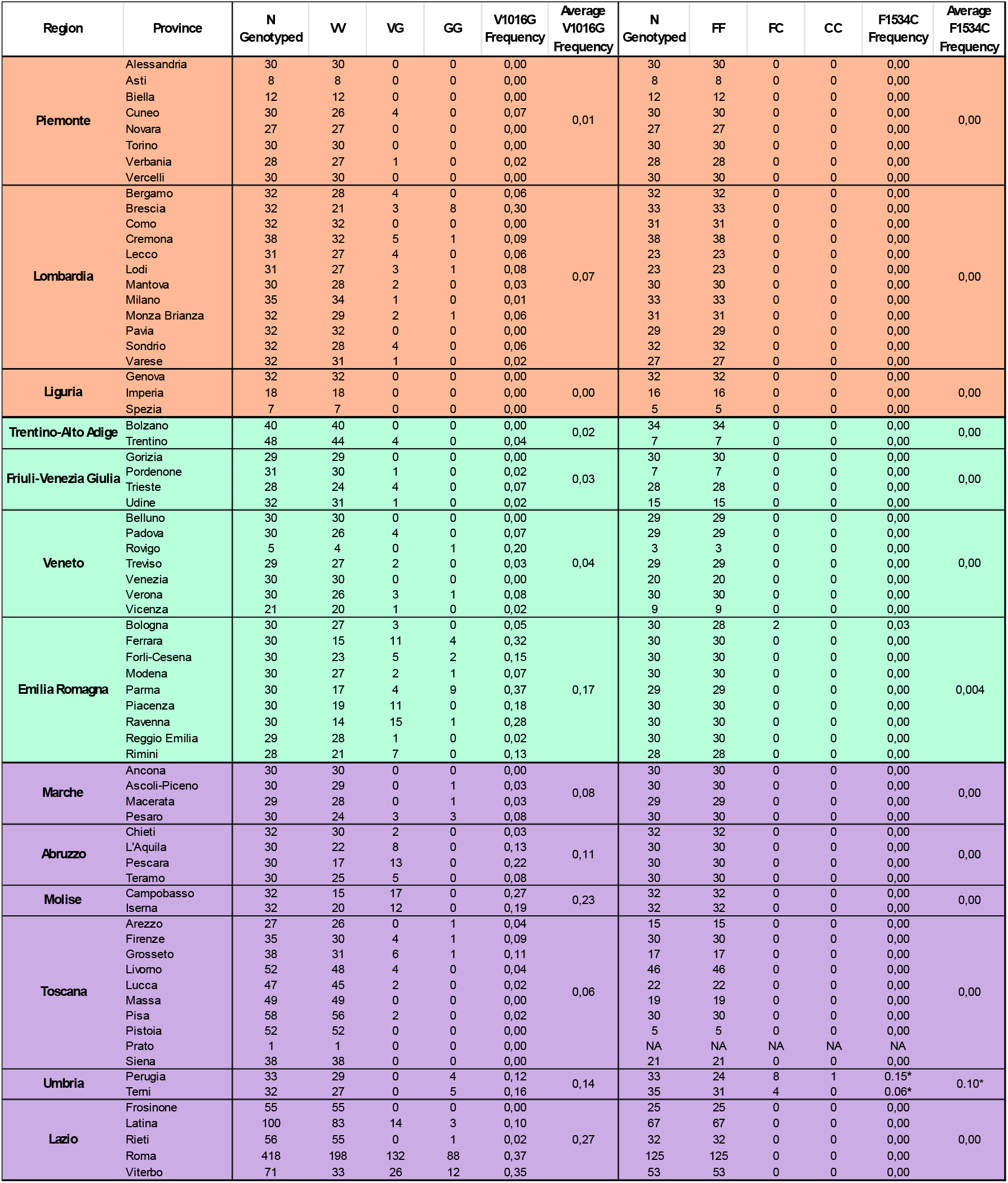
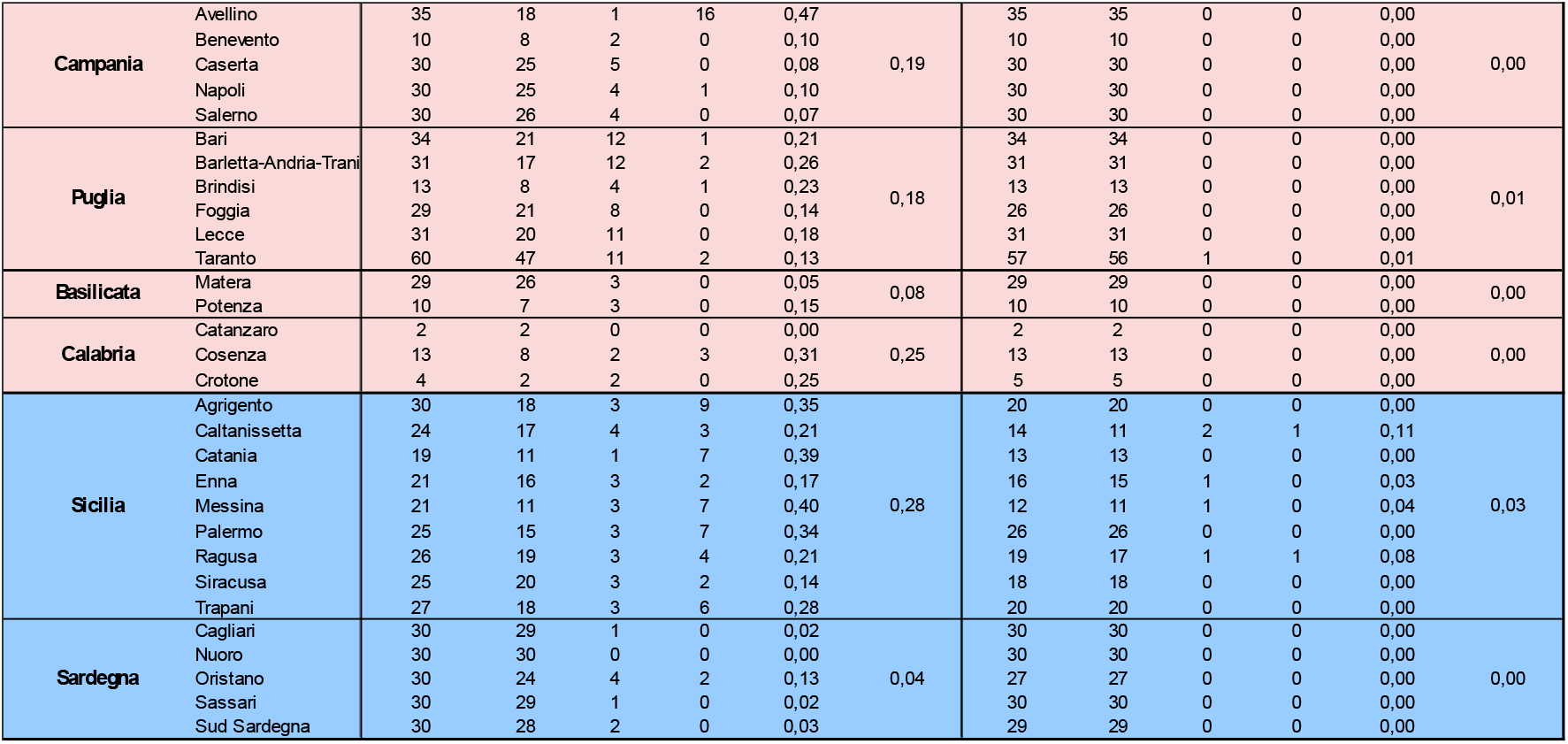
Distribution of *kdr* mutations V1016G and F1534C in *Aedes albopictus* populations collected from various sites across Italian provinces within the MIRIN framework. The table is color-coded according to geographic macro-regions (North-West, North-East, Center, South, and Islands), as depicted in the accompanying Figure 1. NA = Not Available. * FC heterozygous and C/C homozygous genotypes assessed only by PCR (Zhu et al. 2019) and not confirmed by sequencing.

**Figure 1.**
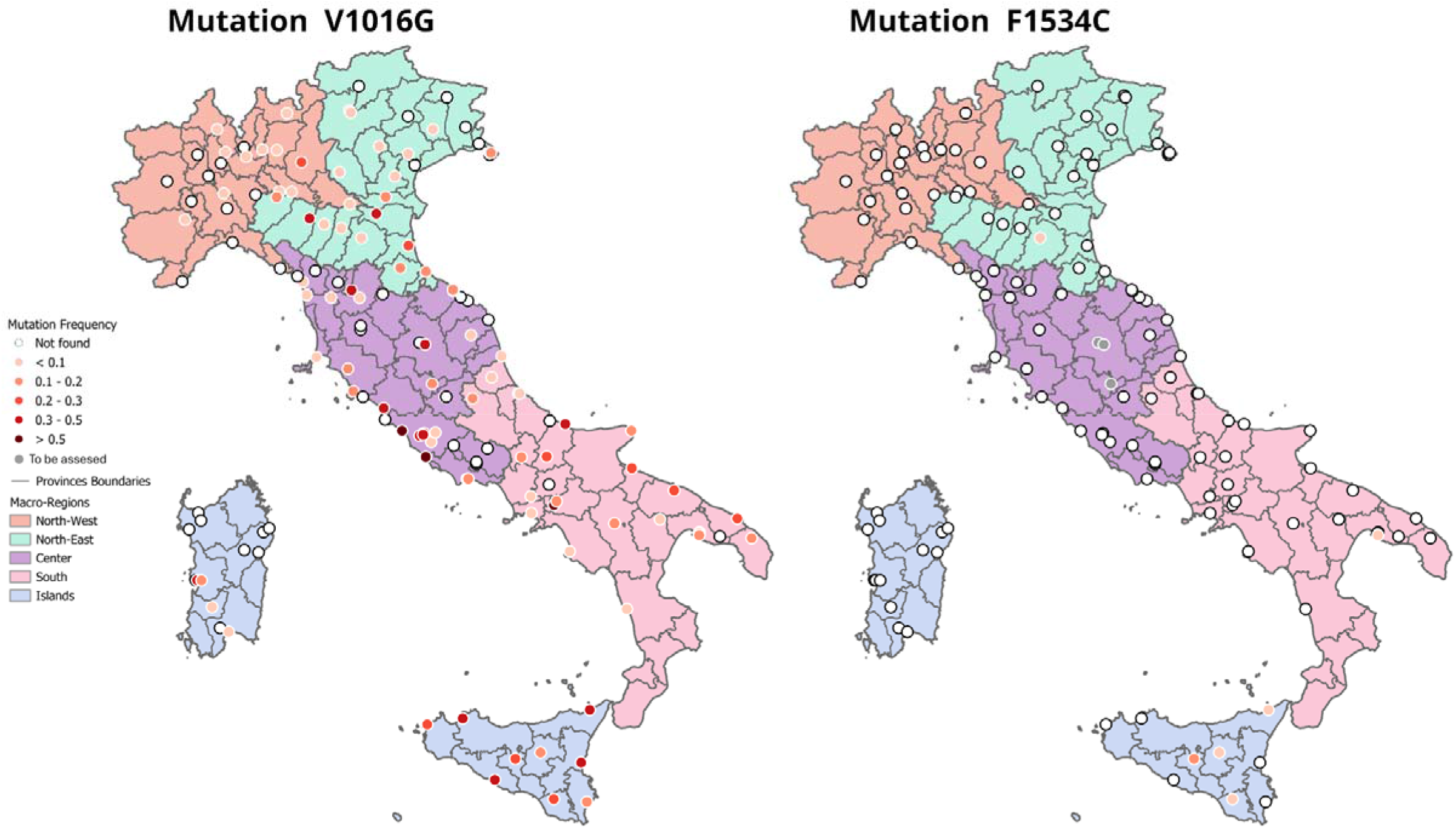
Geographic distribution of *kdr* mutations V1016G (A) and F1534C (B) in *Ae. albopictus* populations across Italy. Each dot represents a sampling site and is or-coded according to the detected mutation frequency (ranging from 0 to >0.5). White dots indicate absence of the mutations. Only sampling sites with at least 5 otyped individuals are shown. The maps are divided into geographic macro-regions (North-West, North-East, Center, South, and Islands) color-coded consistently with le 1. Presence of F1534C in samples from Umbria is highlighted in grey as these have not been confirmed. Provincial boundaries are outlined in grey.

The detailed dataset at site level is available for download from De Marco (2026) in CSV formats. Each row corresponds to a distinct sampling event at a specific location and time point. The dataset captures key information such as sampling coordinates (masked for privacy), collection methods, mosquito life stage, the genotypes and V1016G and F1534C mutation frequencies both at site and level. As described above, to facilitate environmental and spatial analyses, the dataset is enriched with land cover classification, 2m air temperature, and population data aggregated over standardized spatial buffers. All records are structured to ensure consistency and compatibility with geospatial tools. A complete description of each data column of Table 2 is provided below.

**Table 2.**
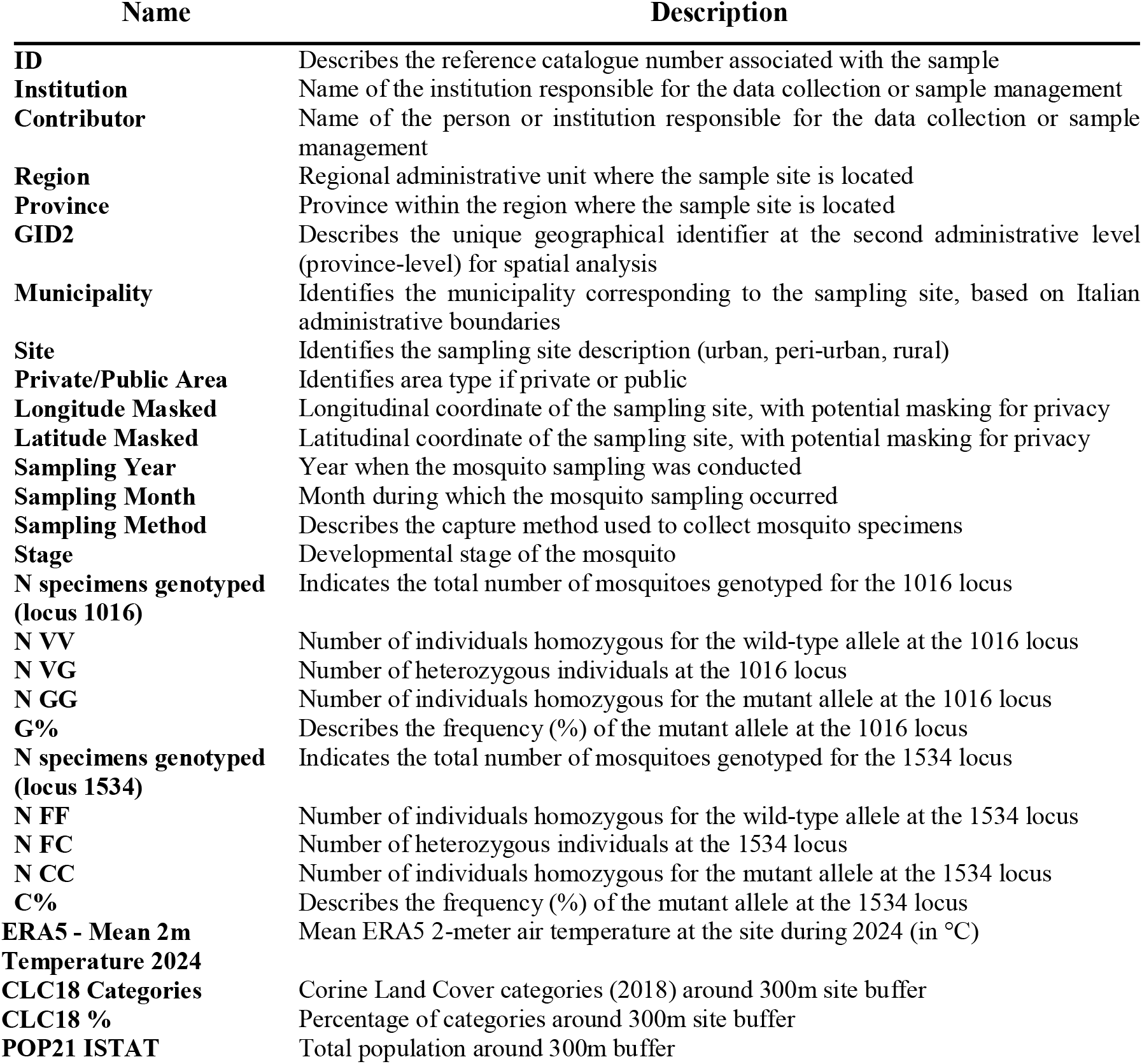
Columns and description of the dataset released. The Table shows the structure of the dataset, listing all columns along with their descriptions. The dataset includes metadata related to the population analysed such as identification, geographical location, sampling methodology, environmental and demographic variables and the genotypic data for two kdr loci (1016 and 1534).

| Name | Description |
| --- | --- |
| **ID** | Describes the reference catalogue number associated with the sample |
| **Institution** | Name of the institution responsible for the data collection or sample management |
| **Contributor** | Name of the person or institution responsible for the data collection or sample management |
| **Region** | Regional administrative unit where the sample site is located |
| **Province** | Province within the region where the sample site is located |
| **GID2** | Describes the unique geographical identifier at the second administrative level (province-level) for spatial analysis |
| **Municipality** | Identifies the municipality corresponding to the sampling site, based on Italian administrative boundaries |
| **Site** | Identifies the sampling site description (urban, peri-urban, rural) |
| **Private/Public Area** | Identifies area type if private or public |
| **Longitude Masked** | Longitudinal coordinate of the sampling site, with potential masking for privacy |
| **Latitude Masked** | Latitudinal coordinate of the sampling site, with potential masking for privacy |
| **Sampling Year** | Year when the mosquito sampling was conducted |
| **Sampling Month** | Month during which the mosquito sampling occurred |
| **Sampling Method** | Describes the capture method used to collect mosquito specimens |
| **Stage** | Developmental stage of the mosquito |
| **N specimens genotyped (locus 1016)** | Indicates the total number of mosquitoes genotyped for the 1016 locus |
| **N VV** | Number of individuals homozygous for the wild-type allele at the 1016 locus |
| **N VG** | Number of heterozygous individuals at the 1016 locus |
| **N GG** | Number of individuals homozygous for the mutant allele at the 1016 locus |
| **G%** | Describes the frequency (%) of the mutant allele at the 1016 locus |
| **N specimens genotyped (locus 1534)** | Indicates the total number of mosquitoes genotyped for the 1534 locus |
| **N FF** | Number of individuals homozygous for the wild-type allele at the 1534 locus |
| **N FC** | Number of heterozygous individuals at the 1534 locus |
| **N CC** | Number of individuals homozygous for the mutant allele at the 1534 locus |
| **C%** | Describes the frequency (%) of the mutant allele at the 1534 locus |
| **ERA5 - Mean 2m Temperature 2024** | Mean ERA5 2-meter air temperature at the site during 2024 (in °C) |
| **CLC18 Categories** | Corine Land Cover categories (2018) around 300m site buffer |
| **CLC18 %** | Percentage of categories around 300m site buffer |
| **POP21 ISTAT** | Total population around 300m buffer |

## TECHNICAL VALIDATION

Collection data were checked by two independent collaborators for internal consistency to ensure that all coordinates for point locations fell in the right nearest centroid of the grid.

Mosquito specimens were identified by experienced taxonomists in each collaborating Institution according with taxonomic keys by Severini et al. (2009).

The genotyping of *Ae. albopictus* specimens for the *kdr* mutations was performed using well standardized and accurate AS-PCR assays, following validated protocols widely used in the scientific community (Pichler et al. 2021 for V1016G mutation; Zhu et al. 2019 for F1534C mutation). All reactions were performed under controlled laboratory conditions in dedicated PCR facilities to minimize cross-contamination. In the rare cases when ambiguous V1016G PCR-banding patterns were obtained the PCR reactions were repeated. Notably, since the F1534C mutation was never observed before in Italy and since the PCR-genotyping protocol is prone to errors (93% accuracy; Pichler et al., 2025), we reassessed the genotype of 25 specimens PCR-genotyped as heterozygous or homozygous for the C allele, either by running the PCR products on a capillary gel (7 specimens from Sicily) or by sequencing (1, 2 and 15 specimens from Liguria, Emilia Romagna and Puglia, respectively). Sequences are available at genbank accession numbers PZ417061–PZ417078. Samples from Umbria were not available for this double-check and are reported in the dataset with an asterisk to highlight that the presence of F1534C mutation was not confirmed.

## REUSE POTENTIAL

We here provide the first set of data at national, regional and province level on the frequencies of the two *kdr* substitutions with strongest effect in conferring pyrethroid resistance in *Ae. albopictus* vector of arbovirus such as Dengue in Italy. It should here be emphasized that pesticide resistance is a dynamic, evolutionary phenomenon and a record in this database may or may not be indicative of actual frequencies in a specific area. Similarly, the absence of a record in this database does not indicate absence of the mutated alleles. Moreover, the data reported are only indicative of target-site resistance, and do not account for other mechanisms such as metabolic resistance.

Nevertheless, this baseline set of data shows that in Italy - which is colonised by *Ae. albopictus* for more than 30 years – the V1016G mutation is widespread in all regions mostly at frequencies <10% per site, but may reach values ranging from 20% to 40% in a few sites. On the other hand, the F1534C mutation was unambiguously detected only in one site in Emilia Romagna, one site in Puglia and in four sites in Sicily, at frequencies ≤3% per site.

According to ECDC (2023) monitoring of biocide resistance in Europe can be improved by establishing links between scientists in biocide resistance research and professionals from the public and veterinary health services involved in vector control. These collaborations could be useful to update biocide resistance monitoring data, generate risk maps, provide scientific and technical expertise to policy makers and disseminate information among stakeholders and countries. This dataset represents a starting point for achieving this goal.

The following potential re-use by researchers, public health officers dealing with mosquito control and resistance management, pest control companies and citizens is envisaged.

### Scientific community

researchers can exploit genotypic, geographic and ecological data (and possibly enrich the database with additional information, including historical and future genotypic data) in order to carry out specific analyses. For example, data can be exploited to understand the spatial and temporal dynamics of spreading of the mutated alleles in Italy and generate risk maps.

### Public health officers planning mosquito control

the availability of baseline data and risk maps on the spatial distribution of alleles associated with pyrethroid resistance will facilitate the identification of possible resistant *Ae. albopictus* populations at site/province level, where bioassays should be carried out to assess for phenotypic resistance to pyrethroids, as requested by the PNA 2020-2025 (Ministero della Salute, 2019). These populations could eventually be used as “sentinels” to be monitored over time. Based on this, the public health administration may plan evidence-based mosquito control and insecticide resistance management.

<u>Pest control companies</u> may exploit the dataset to predict where a higher risk of failure of pyrethroid treatments exists, for instance in site/provinces where both V1016G and F1534C alleles coexist and may synergise increasing the resistant phenotype, as shown to occur in the tropical arbovirus vector species, *Ae. aegypti* (Hirata et al., 2014).

<u>Citizen</u>, in addition to data on spreading of pyrethroid resistance in Italy, may easily access information on sites/provinces where selective pressure likely associated with excessive use of pyrethroids is highest and have the possibility to actively and knowingly influence mosquito control planning in private and public areas.

## Availability of Source Code and Requirements

Project name: irdash

Project homepage: https://github.com/randomxsk8/irdash

Operating system: Linux/ Windows/macOS Programming language: Python

Other requirements: pandas, geopandas, shapely, cdsapi, xarray

License: MIT license

## ACKNOWLEDGEMENTS

This work received financial support from Ministero dell’Università e della Ricerca (MUR) in the framework of EU funding within the Next Generation EU MUR PNRR Extended Partnership initiative on Emerging Infectious Diseases (Project no. PE00000007, INF-ACT).

## REFERENCES

Bossche, J. Van den, Jordahl, K., Fleischmann, M., Richards, M., McBride, J., Wasserman, J., Badaracco, A.G., Snow, A.D., Roggemans, P., Ward, B., Tratner, J., Gerard, J., Perry, M., Taves, M., carsonfarmer, Hjelle G.A., Bell, R. Hoeven, E. ter, Cochran, M., Tan, N.Y., rraymondgh, Caria G., Culbertson, L., Bartos, M., Rey, S., Flavin, J., Eubank, N., sangarshanan, Gillies S., 2025. geopandas/geopandas: v1.1.1. 10.5281/ZENODO.15750510

Dash Enterprise. Plotly Technologies Inc., 2025. Dash Documentation & User Guide | Plotly [WWW Document]. URL https://dash.plotly.com/ (accessed 7.31.25).

De Marco CM, Schaffner F (2026). Systematic literature review on the distribution of priority mosquito species within the VectorNet geographical area. European Food Safety Authority. Occurrence dataset 10.15468/kvat4a

De Marco, C. M. (2026). irdash (Version 1.0.0) [Computer software]. https://github.com/randomxsk8/irdash

ECDC, 2025a. Historical data on local transmission of dengue in the EU/EEA [WWW Document]. URL https://www.ecdc.europa.eu/en/all-topics-z/dengue/surveillance-and-disease-data/autochthonous-transmission-dengue-virus-eueea-previous-years

ECDC, 2025b. Seasonal surveillance for chikungunya virus disease in the EU/EEA for 2025 [WWW Document]. URL https://www.ecdc.europa.eu/en/chikungunya-virus-disease/surveillance-and-updates/seasonal-surveillance

ECDC, 2024. Worsening spread of mosquito-borne disease outbreaks in EU/EEA, according to latest ECDC figures [WWW Document]. URL https://www.ecdc.europa.eu/en/news-events/worsening-spread-mosquito-borne-disease-outbreaks-eueea-according-latest-ecdc-figures (accessed 7.31.25).

ECDC, 2023. Literature review on the state of biocide resistance in wild vector populations in the EU and neighbouring countries.

ECDC, 2012. Guidelines for mosquito surveillance [WWW Document]. URL https://www.ecdc.europa.eu/en/disease-vectors/surveillance-and-disease-data/guidelines-mosquito (accessed 7.31.25).

GADM, 2024. Global ADMinistrative Areas (GADM) [WWW Document]. URL https://gadm.org/index.html

Hemingway, J., Hawkes, N.J., McCarroll, L., Ranson, H., 2004. The molecular basis of insecticide resistance in mosquitoes. Insect Biochem Mol Biol 34, 653–665. 10.1016/j.ibmb.2004.03.018

Hirata, K., Komagata, O., Itokawa, K., Yamamoto, A., Tomita, T., Kasai, S., 2014. A Single Crossing-Over Event in Voltage-Sensitive Na+ Channel Genes May Cause Critical Failure of Dengue Mosquito Control by Insecticides. PLoS Negl Trop Dis 8, e3085. 10.1371/JOURNAL.PNTD.0003085

ISPRA, 2018. Corine Land Cover [WWW Document]. URL https://www.isprambiente.gov.it/it/attivita/suolo-e-territorio/suolo/copertura-del-suolo/corine-land-cover (accessed 7.31.25).

ISTAT, 2021. Basi territoriali e variabili censuarie – Istat [WWW Document]. URL https://www.istat.it/notizia/basi-territoriali-e-variabili-censuarie/ (accessed 7.31.25).

Kasai, S., Caputo, B., Tsunoda, T., Cuong, T.C., Maekaw, Y., Lam-Phua, S.G., Pichler, V., Itokawa, K., Murota, K., Komagata, O., Yoshida, C., Chung, H.H., Bellini, R., Tsuda, Y., Teng, H.J., Lima Filho, J.L. de, Alves, L.C., Ng, L.C., Minakawa, N., Yen, N.T., Phong, T.V., Sawabe, K., Tomita, T., 2019. First detection of a Vssc allele V1016G conferring a high level of insecticide resistance in Aedes albopictus collected from Europe (Italy) and Asia (Vietnam), 2016: A new emerging threat to controlling arboviral diseases. Eurosurveillance 24. 10.2807/1560-7917.ES.2019.24.5.1700847,

Ministero della Salute, 2019. Piano nazionale di prevenzione, sorveglianza e risposta alle Arbovirosi (PNA) 2020-2025 [WWW Document]. URL https://www.salute.gov.it/portale/documentazione/p6_2_2_1.jsp?id=2947 (accessed 8.5.25).

Moyes, C.L., Vontas, J., Martins, A.J., Ng, L.C., Koou, S.Y., Dusfour, I., Raghavendra, K., Pinto, J., Corbel, V., David, J.P., Weetman, D., 2017. Contemporary status of insecticide resistance in the major Aedes vectors of arboviruses infecting humans. PLoS Negl Trop Dis 11. 10.1371/JOURNAL.PNTD.0005625

Muñoz Sabater, J., 2019. ERA5-Land hourly data from 1950 to present. Copernicus Climate Change Service (C3S) Climate Data Store (CDS) [WWW Document]. 10.24381/cds.e2161bac

Pichler V, Valadas V, Akiner MM, Balatsos G, Barceló C, Borg ML, Bouyer J, Bravo-Barriga D, Bueno R, Caputo B, Collantes F, Delacour-Estrella S, Velo E, Falcuta E, Flacio E, García-Pérez AL, Gómez JF, Horvath C, Adam K, Kadriaj P, Kavran M, L’Ambert G, Lia RP, Marabuto E, Medialdea-Carrera R, Melero-Alcibar R, Michaelakis A, Mihalca AD, Micocci M, Mikov O, Miranda MA, Müller P, Ornosa C, Outerelo R, Otranto D, Pajovic I, Pérez-Tris J, Petric D, Rebelo MT, Besnard G, Rogozi E, Tello A, Vázquez Á, Vasquez M, Zitko T, Schaffner F, Della Torre A, Pinto J. Tracking pyrethroid resistance in arbovirus mosquito vectors: mutations I1532T and F1534C in Aedes albopictus across Europe. Parasit Vectors. 2025 Dec 24;18(1):506. doi: 10.1186/s13071-025-07130-1. PMID: 41444593; PMCID: PMC12728990.

Pichler, V., Caputo, B., Valadas, V., Micocci, M., Horvath, C., Virgillito, C., Akiner, M., Balatsos, G., Bender, C., Besnard, G., Bravo-Barriga, D., Bueno-Mari, R., Collantes, F., Delacour-Estrella, S., Dikolli, E., Falcuta, E., Flacio, E., García-Pérez, A.L., Kalan, K., Kavran, M., L’Ambert, G., Lia, R.P., Marabuto, E., Medialdea, R., Melero-Alcibar, R., Michaelakis, A., Mihalca, A., Mikov, O., Miranda, M.A., Müller, P., Otranto, D., Pajovic, I., Petric, D., Rebelo, M.T., Robert, V., Rogozi, E., Tello, A., Zitko, T., Schaffner, F., Pinto, J., della Torre, A., 2022. Geographic distribution of the V1016G knockdown resistance mutation in Aedes albopictus: a warning bell for Europe. Parasit Vectors 15, 280. 10.1186/S13071-022-05407-3

Pichler, V., Mancini, E., Micocci, M., Calzetta, M., Arnoldi, D., Rizzoli, A., Lencioni, V., Paoli, F., Bellini, R., Veronesi, R., Martini, S., Drago, A., De Liberato, C., Ermenegildi, A., Pinto, J., Torre, A. Della, Caputo, B., 2021. A novel allele specific polymerase chain reaction (As-pcr) assay to detect the v1016g knockdown resistance mutation confirms its widespread presence in aedes albopictus populations from italy. Insects 12, 1–12. 10.3390/INSECTS12010079

Pichler V, Malandruccolo C, Serini P, Bellini R, Severini F, Toma L, Di Luca M, Montarsi F, Ballardini M, Manica M, Petrarca V, Vontas J, Kasai S, Della Torre A, Caputo B. Phenotypic and genotypic pyrethroid resistance of Aedes albopictus, with focus on the 2017 chikungunya outbreak in Italy. Pest Manag Sci. 2019 Oct;75(10):2642–2651. doi: 10.1002/ps.5369. Epub 2019 Apr 8. PMID: 30729706.

Pichler V, Bellini R, Veronesi R, Arnoldi D, Rizzoli A, Lia RP, Otranto D, Montarsi F, Carlin S, Ballardini M, Antognini E, Salvemini M, Brianti E, Gaglio G, Manica M, Cobre P, Serini P, Velo E, Vontas J, Kioulos I, Pinto J, Della Torre A, Caputo B. First evidence of resistance to pyrethroid insecticides in Italian Aedes albopictus populations 26 years after invasion. Pest Manag Sci. 2018 Jun;74(6):1319–1327. doi: 10.1002/ps.4840. Epub 2018 Feb 21. PMID: 29278457.

Plotly Technologies Inc., 2025. Plotly Python Graphing Library [WWW Document]. URL https://plotly.com/python/ (accessed 7.31.25).

Schaffner, F., Medlock, J.M., Van Bortel, W., 2013. Public health significance of invasive mosquitoes in Europe. Clinical Microbiology and Infection 19, 685–692. 10.1111/1469-0691.12189

Severini, F., Toma, L., Luca, M. Di, Romi, R., 2009. Le zanzare italiane: generalità e identificazione degli adulti (diptera, Culicidae). Fragm Entomol 41, 213–372. 10.13133/2284-4880/92

Uemura, N., Itokawa, K., Komagata, O., Kasai, S., 2024. Recent advances in the study of knockdown resistance mutations in Aedes mosquitoes with a focus on several remarkable mutations. Curr Opin Insect Sci 63, 101178. 10.1016/j.cois.2024.101178

Zhu, C.Y., Zhao, C.C., Wang, Y.G., Ma, D.L., Song, X.P., Wang, J., Meng, F.X., 2019. Establishment of an innovative and sustainable PCR technique for 1534 locus mutation of the knockdown resistance (kdr) gene in the dengue vector Aedes albopictus. Parasit Vectors 12, 1–8. 10.1186/S13071-019-3829-5

