## Supplementary material for "Tracking *kdr* Alleles Associated with Pyrethroid Resistance in *Aedes albopictus* across Italy: A Nationwide Genotypic Dataset by MosqIRIT Network": Table 1: Table_1.pdf

| Region | Province | N Genotyped | VV | VG | GG | V1016G Frequency | Average V1016G Frequency | N Genotyped | FF | FC | CC | F1534C Frequency | Average F1534C Frequency |
|---|---|---|---|---|---|---|---|---|---|---|---|---|---|
| **Piemonte** | Alessandria | 30 | 30 | 0 | 0 | 0,00 | 0,01 | 30 | 30 | 0 | 0 | 0,00 | 0,00 |
| | Asti | 8 | 8 | 0 | 0 | 0,00 | | 8 | 8 | 0 | 0 | 0,00 | |
| | Biella | 12 | 12 | 0 | 0 | 0,00 | | 12 | 12 | 0 | 0 | 0,00 | |
| | Cuneo | 30 | 26 | 4 | 0 | 0,07 | | 30 | 30 | 0 | 0 | 0,00 | |
| | Novara | 27 | 27 | 0 | 0 | 0,00 | | 27 | 27 | 0 | 0 | 0,00 | |
| | Torino | 30 | 30 | 0 | 0 | 0,00 | | 30 | 30 | 0 | 0 | 0,00 | |
| | Verbania | 28 | 27 | 1 | 0 | 0,02 | | 28 | 28 | 0 | 0 | 0,00 | |
| | Vercelli | 30 | 30 | 0 | 0 | 0,00 | | 30 | 30 | 0 | 0 | 0,00 | |
| **Lombardia** | Bergamo | 32 | 28 | 4 | 0 | 0,06 | 0,07 | 32 | 32 | 0 | 0 | 0,00 | 0,00 |
| | Brescia | 32 | 21 | 3 | 8 | 0,30 | | 33 | 33 | 0 | 0 | 0,00 | |
| | Como | 32 | 32 | 0 | 0 | 0,00 | | 31 | 31 | 0 | 0 | 0,00 | |
| | Cremona | 38 | 32 | 5 | 1 | 0,09 | | 38 | 38 | 0 | 0 | 0,00 | |
| | Lecco | 31 | 27 | 4 | 0 | 0,06 | | 23 | 23 | 0 | 0 | 0,00 | |
| | Lodi | 31 | 27 | 3 | 1 | 0,08 | | 23 | 23 | 0 | 0 | 0,00 | |
| | Mantova | 30 | 28 | 2 | 0 | 0,03 | | 30 | 30 | 0 | 0 | 0,00 | |
| | Milano | 35 | 34 | 1 | 0 | 0,01 | | 33 | 33 | 0 | 0 | 0,00 | |
| | Monza Brianza | 32 | 29 | 2 | 1 | 0,06 | | 31 | 31 | 0 | 0 | 0,00 | |
| | Pavia | 32 | 32 | 0 | 0 | 0,00 | | 29 | 29 | 0 | 0 | 0,00 | |
| | Sondrio | 32 | 28 | 4 | 0 | 0,06 | | 32 | 32 | 0 | 0 | 0,00 | |
| | Varese | 32 | 31 | 1 | 0 | 0,02 | | 27 | 27 | 0 | 0 | 0,00 | |
| **Liguria** | Genova | 32 | 32 | 0 | 0 | 0,00 | 0,00 | 32 | 32 | 0 | 0 | 0,00 | 0,00 |
| | Imperia | 18 | 18 | 0 | 0 | 0,00 | | 16 | 16 | 0 | 0 | 0,00 | |
| | Spezia | 7 | 7 | 0 | 0 | 0,00 | | 5 | 5 | 0 | 0 | 0,00 | |
| **Trentino-Alto Adige** | Bolzano | 40 | 40 | 0 | 0 | 0,00 | 0,02 | 34 | 34 | 0 | 0 | 0,00 | 0,00 |
| | Trentino | 48 | 44 | 4 | 0 | 0,04 | | 7 | 7 | 0 | 0 | 0,00 | |
| **Friuli-Venezia Giulia** | Gorizia | 29 | 29 | 0 | 0 | 0,00 | 0,03 | 30 | 30 | 0 | 0 | 0,00 | 0,00 |
| | Pordenone | 31 | 30 | 1 | 0 | 0,02 | | 7 | 7 | 0 | 0 | 0,00 | |
| | Trieste | 28 | 24 | 4 | 0 | 0,07 | | 28 | 28 | 0 | 0 | 0,00 | |
| | Udine | 32 | 31 | 1 | 0 | 0,02 | | 15 | 15 | 0 | 0 | 0,00 | |
| **Veneto** | Belluno | 30 | 30 | 0 | 0 | 0,00 | 0,04 | 29 | 29 | 0 | 0 | 0,00 | 0,00 |
| | Padova | 30 | 26 | 4 | 0 | 0,07 | | 29 | 29 | 0 | 0 | 0,00 | |
| | Rovigo | 5 | 4 | 0 | 1 | 0,20 | | 3 | 3 | 0 | 0 | 0,00 | |
| | Treviso | 29 | 27 | 2 | 0 | 0,03 | | 29 | 29 | 0 | 0 | 0,00 | |
| | Venezia | 30 | 30 | 0 | 0 | 0,00 | | 20 | 20 | 0 | 0 | 0,00 | |
| | Verona | 30 | 26 | 3 | 1 | 0,08 | | 30 | 30 | 0 | 0 | 0,00 | |
| | Vicenza | 21 | 20 | 1 | 0 | 0,02 | | 9 | 9 | 0 | 0 | 0,00 | |
| **Emilia Romagna** | Bologna | 30 | 27 | 3 | 0 | 0,05 | 0,17 | 30 | 28 | 2 | 0 | 0,03 | 0,004 |
| | Ferrara | 30 | 15 | 11 | 4 | 0,32 | | 30 | 30 | 0 | 0 | 0,00 | |
| | Forlì-Cesena | 30 | 23 | 5 | 2 | 0,15 | | 30 | 30 | 0 | 0 | 0,00 | |
| | Modena | 30 | 27 | 2 | 1 | 0,07 | | 30 | 30 | 0 | 0 | 0,00 | |
| | Parma | 30 | 17 | 4 | 9 | 0,37 | | 29 | 29 | 0 | 0 | 0,00 | |
| | Piacenza | 30 | 19 | 11 | 0 | 0,18 | | 30 | 30 | 0 | 0 | 0,00 | |
| | Ravenna | 30 | 14 | 15 | 1 | 0,28 | | 30 | 30 | 0 | 0 | 0,00 | |
| | Reggio Emilia | 29 | 28 | 1 | 0 | 0,02 | | 30 | 30 | 0 | 0 | 0,00 | |
| | Rimini | 28 | 21 | 7 | 0 | 0,13 | | 28 | 28 | 0 | 0 | 0,00 | |
| **Marche** | Ancona | 30 | 30 | 0 | 0 | 0,00 | 0,08 | 30 | 30 | 0 | 0 | 0,00 | 0,00 |
| | Ascoli-Piceno | 30 | 29 | 0 | 1 | 0,03 | | 30 | 30 | 0 | 0 | 0,00 | |
| | Macerata | 29 | 28 | 0 | 1 | 0,03 | | 29 | 29 | 0 | 0 | 0,00 | |
| | Pesaro | 30 | 24 | 3 | 3 | 0,08 | | 30 | 30 | 0 | 0 | 0,00 | |
| **Abruzzo** | Chieti | 32 | 30 | 2 | 0 | 0,03 | 0,11 | 32 | 32 | 0 | 0 | 0,00 | 0,00 |
| | L'Aquila | 30 | 22 | 8 | 0 | 0,13 | | 30 | 30 | 0 | 0 | 0,00 | |
| | Pescara | 30 | 17 | 13 | 0 | 0,22 | | 30 | 30 | 0 | 0 | 0,00 | |
| | Teramo | 30 | 25 | 5 | 0 | 0,08 | | 30 | 30 | 0 | 0 | 0,00 | |
| **Molise** | Campobasso | 32 | 15 | 17 | 0 | 0,27 | 0,23 | 32 | 32 | 0 | 0 | 0,00 | 0,00 |
| | Iserna | 32 | 20 | 12 | 0 | 0,19 | | 32 | 32 | 0 | 0 | 0,00 | |
| **Toscana** | Arezzo | 27 | 26 | 0 | 1 | 0,04 | 0,06 | 15 | 15 | 0 | 0 | 0,00 | 0,00 |
| | Firenze | 35 | 30 | 4 | 1 | 0,09 | | 30 | 30 | 0 | 0 | 0,00 | |
| | Grosseto | 38 | 31 | 6 | 1 | 0,11 | | 17 | 17 | 0 | 0 | 0,00 | |
| | Livorno | 52 | 48 | 4 | 0 | 0,04 | | 46 | 46 | 0 | 0 | 0,00 | |
| | Lucca | 47 | 45 | 2 | 0 | 0,02 | | 22 | 22 | 0 | 0 | 0,00 | |
| | Massa | 49 | 49 | 0 | 0 | 0,00 | | 19 | 19 | 0 | 0 | 0,00 | |
| | Pisa | 58 | 56 | 2 | 0 | 0,02 | | 30 | 30 | 0 | 0 | 0,00 | |
| | Pistoia | 52 | 52 | 0 | 0 | 0,00 | | 5 | 5 | 0 | 0 | 0,00 | |
| | Prato | 1 | 1 | 0 | 0 | 0,00 | | NA | NA | NA | NA | NA | |
| | Siena | 38 | 38 | 0 | 0 | 0,00 | | 21 | 21 | 0 | 0 | 0,00 | |
| **Umbria** | Perugia | 33 | 29 | 0 | 4 | 0,12 | 0,14 | 33 | 24 | 8 | 1 | 0.15* | 0.10* |
| | Terni | 32 | 27 | 0 | 5 | 0,16 | | 35 | 31 | 4 | 0 | 0.06* | |
| **Lazio** | Frosinone | 55 | 55 | 0 | 0 | 0,00 | 0,27 | 25 | 25 | 0 | 0 | 0,00 | 0,00 |
| | Latina | 100 | 83 | 14 | 3 | 0,10 | | 67 | 67 | 0 | 0 | 0,00 | |
| | Rieti | 56 | 55 | 0 | 1 | 0,02 | | 32 | 32 | 0 | 0 | 0,00 | |
| | Roma | 418 | 198 | 132 | 88 | 0,37 | | 125 | 125 | 0 | 0 | 0,00 | |
| | Viterbo | 71 | 33 | 26 | 12 | 0,35 | | 53 | 53 | 0 | 0 | 0,00 | |
| **Campania** | Avellino | 35 | 18 | 1 | 16 | 0,47 | 0,19 | 35 | 35 | 0 | 0 | 0,00 | 0,00 |
| | Benevento | 10 | 8 | 2 | 0 | 0,10 | | 10 | 10 | 0 | 0 | 0,00 | |
| | Caserta | 30 | 25 | 5 | 0 | 0,08 | | 30 | 30 | 0 | 0 | 0,00 | |
| | Napoli | 30 | 25 | 4 | 1 | 0,10 | | 30 | 30 | 0 | 0 | 0,00 | |
| | Salerno | 30 | 26 | 4 | 0 | 0,07 | | 30 | 30 | 0 | 0 | 0,00 | |
| **Puglia** | Bari | 34 | 21 | 12 | 1 | 0,21 | 0,18 | 34 | 34 | 0 | 0 | 0,00 | 0,01 |
| | Barletta-Andria-Trani | 31 | 17 | 12 | 2 | 0,26 | | 31 | 31 | 0 | 0 | 0,00 | |
| | Brindisi | 13 | 8 | 4 | 1 | 0,23 | | 13 | 13 | 0 | 0 | 0,00 | |
| | Foggia | 29 | 21 | 8 | 0 | 0,14 | | 26 | 26 | 0 | 0 | 0,00 | |
| | Lecce | 31 | 20 | 11 | 0 | 0,18 | | 31 | 31 | 0 | 0 | 0,00 | |
| | Taranto | 60 | 47 | 11 | 2 | 0,13 | | 57 | 56 | 1 | 0 | 0,01 | |
| **Basilicata** | Matera | 29 | 26 | 3 | 0 | 0,05 | 0,08 | 29 | 29 | 0 | 0 | 0,00 | 0,00 |
| | Potenza | 10 | 7 | 3 | 0 | 0,15 | | 10 | 10 | 0 | 0 | 0,00 | |
| **Calabria** | Catanzaro | 2 | 2 | 0 | 0 | 0,00 | 0,25 | 2 | 2 | 0 | 0 | 0,00 | 0,00 |
| | Cosenza | 13 | 8 | 2 | 3 | 0,31 | | 13 | 13 | 0 | 0 | 0,00 | |
| | Crotone | 4 | 2 | 2 | 0 | 0,25 | | 5 | 5 | 0 | 0 | 0,00 | |
| **Sicilia** | Agrigento | 30 | 18 | 3 | 9 | 0,35 | 0,28 | 20 | 20 | 0 | 0 | 0,00 | 0,03 |
| | Caltanissetta | 24 | 17 | 4 | 3 | 0,21 | | 14 | 11 | 2 | 1 | 0,11 | |
| | Catania | 19 | 11 | 1 | 7 | 0,39 | | 13 | 13 | 0 | 0 | 0,00 | |
| | Enna | 21 | 16 | 3 | 2 | 0,17 | | 16 | 15 | 1 | 0 | 0,03 | |
| | Messina | 21 | 11 | 3 | 7 | 0,40 | | 12 | 11 | 1 | 0 | 0,04 | |
| | Palermo | 25 | 15 | 3 | 7 | 0,34 | | 26 | 26 | 0 | 0 | 0,00 | |
| | Ragusa | 26 | 19 | 3 | 4 | 0,21 | | 19 | 17 | 1 | 1 | 0,08 | |
| | Siracusa | 25 | 20 | 3 | 2 | 0,14 | | 18 | 18 | 0 | 0 | 0,00 | |
| | Trapani | 27 | 18 | 3 | 6 | 0,28 | | 20 | 20 | 0 | 0 | 0,00 | |
| **Sardegna** | Cagliari | 30 | 29 | 1 | 0 | 0,02 | 0,04 | 30 | 30 | 0 | 0 | 0,00 | 0,00 |
| | Nuoro | 30 | 30 | 0 | 0 | 0,00 | | 30 | 30 | 0 | 0 | 0,00 | |
| | Oristano | 30 | 24 | 4 | 2 | 0,13 | | 27 | 27 | 0 | 0 | 0,00 | |
| | Sassari | 30 | 29 | 1 | 0 | 0,02 | | 30 | 30 | 0 | 0 | 0,00 | |
| | Sud Sardegna | 30 | 28 | 2 | 0 | 0,03 | | 29 | 29 | 0 | 0 | 0,00 | |
